# HiC-TE: a computational pipeline for Hi-C data analysis shows a possible role of repeat family interactions in the genome 3D organization

**DOI:** 10.1101/2021.12.18.473300

**Authors:** Matej Lexa, Monika Cechova, Son Hoang Nguyen, Pavel Jedlicka, Viktor Tokan, Zdenek Kubat, Roman Hobza, Eduard Kejnovsky

## Abstract

The role of repetitive DNA in the 3D organization of the interphase nucleus in plant cells is a subject of intensive study. High-throughput chromosome conformation capture (Hi-C) is a sequencing-based method detecting the proximity of DNA segments in nuclei. We combined Hi-C data, plant reference genome data and tools for the characterization of genomic repeats to build a Nextflow pipeline identifying and quantifying the contacts of specific repeats revealing the preferential homotypic interactions of ribosomal DNA, DNA transposons and some LTR retrotransposon families. We provide a novel way to analyze the organization of repetitive elements in the 3D nucleus.

## Background

A significant part of eukaryotic genomes, namely in plants, comprises transposable elements (TEs) and satellite DNA, where e.g. LTR retrotransposons constitute up to 90% of genomes [1–3]. Despite their initial consideration as junk DNA, many functions of repeats have been revealed during the last decades demonstrating their role in the structure and function of genomes and cells. TEs are often embedded in cellular regulatory networks [4] where they re-wire the gene expression programs [5]. Many examples of the domestication of TEs for specific cellular functions have been observed [6, 7]. The eukaryotic genome is hierarchically packed in the nucleus allowing DNA replication and gene transcription to take place in a spatially and temporally regulated fashion. Distinct organizations of chromosomes in interphase nuclei were revealed exhibiting interactions of centromeric or telomeric repetitive DNA.

Methods of high-throughput mapping of DNA-DNA interactions, such as chromosome conformation capture (Hi-C), now allow the study of long-distance interactions in nuclei. A better understanding of the interaction of the main repeat classes can help uncover their genomic role. A recent study demonstrated the role of TEs in organizing the human and mouse genomes [8] but similar analysis in plants is hitherto missing. Additionally, since centromeres and telomeres are mostly composed of repetitive DNA, such analyses have the potential to verify the Rabl and Rosette organizations at the molecular level.

Here we present a new sequence processing pipeline to identify and quantify interactions of transposable elements, satellite DNA and rDNA in nuclei, especially those that participate in long-distance (>1Mbp) or interchromosomal contacts with frequencies that differ from the baseline expectations.

## Results

Our pipeline (Fig.1, Additional File 1) integrates genome sequence analysis from several sources: assembled genome annotation for the transposable elements and satellite DNA (TE-greedy-nester, PlantSat database), medium and long-distance contact information (Hi-C sequencing experiments) and repetitive sequencing read clustering (Repeat Explorer 2). The pipeline was implemented with Nextflow to allow for flexibility and scalability, using a recent installation of Ubuntu Linux with all dependencies included. In addition, we provide a tested containerized version allowing runs with Docker/Singularity deployment (manual and alternative test config file provided, see Additional File 2 and pipeline repository). As a result, all the figures and tables are fully reproducible and can be easily generated. We summarize the memory, disk, and time requirements in Additional File 3 and 4.

**Figure 1.**
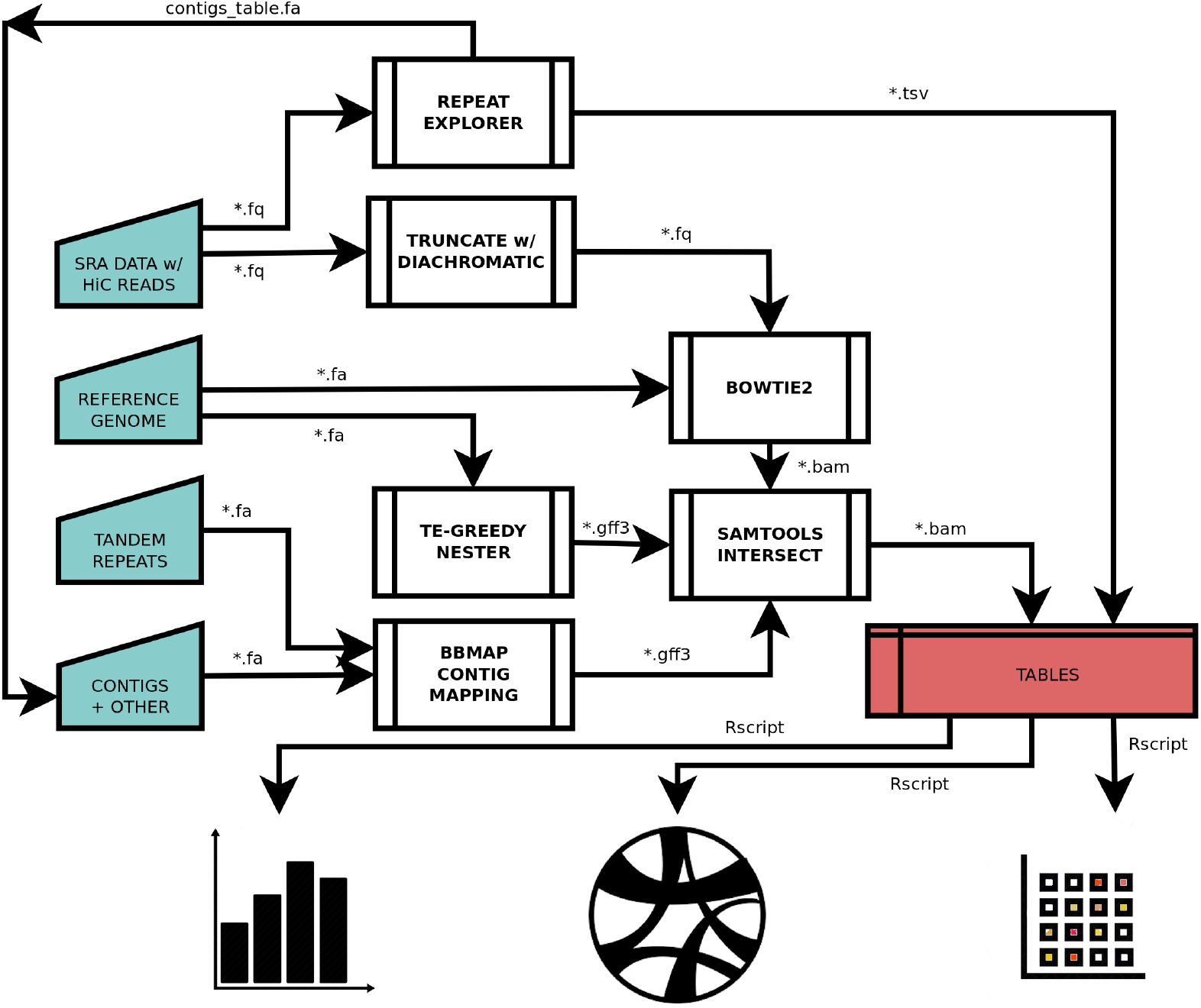
Block diagram showing the overall data flow in the HiC-TE pipeline. Some details were omitted for greater clarity (the full graph as produced by the pipeline is shown in Additional File 1)(blue - main data inputs; red - main data outputs; double edged rectangles - main processes running external bioinformatics tools; FASTA (^*^.fa), FASTQ (^*^.fq), BAM, GFF3 - main sequence and annotation data formats passed between processes; Rscript - R visualization scripts).

To test the pipeline, we used a publicly available dataset with six independent Hi-C experiments on the tomato (Solanum lycopersicum) with two technical replicates for each of three plants (from [9]. We verified that the pipeline produces consistent results and that the computational replicates are less variable than any other replicates present. We analyzed Hi-C contacts from reads clustered with Repeat Explorer (Fig.2a) and reference-mapped long-distance interactions (spanning more than 1 Mbp or between sequences located on different chromosomes)(Fig.2b). The main output is a series of heatmaps showing high and low values of normalized contacts in diverging colors, while fields (repeat family pairs) with missing values are shown in grey. This is typically caused by extremely low copy number of one of the families (either in data or after normalization randomization).

**Figure 2.**
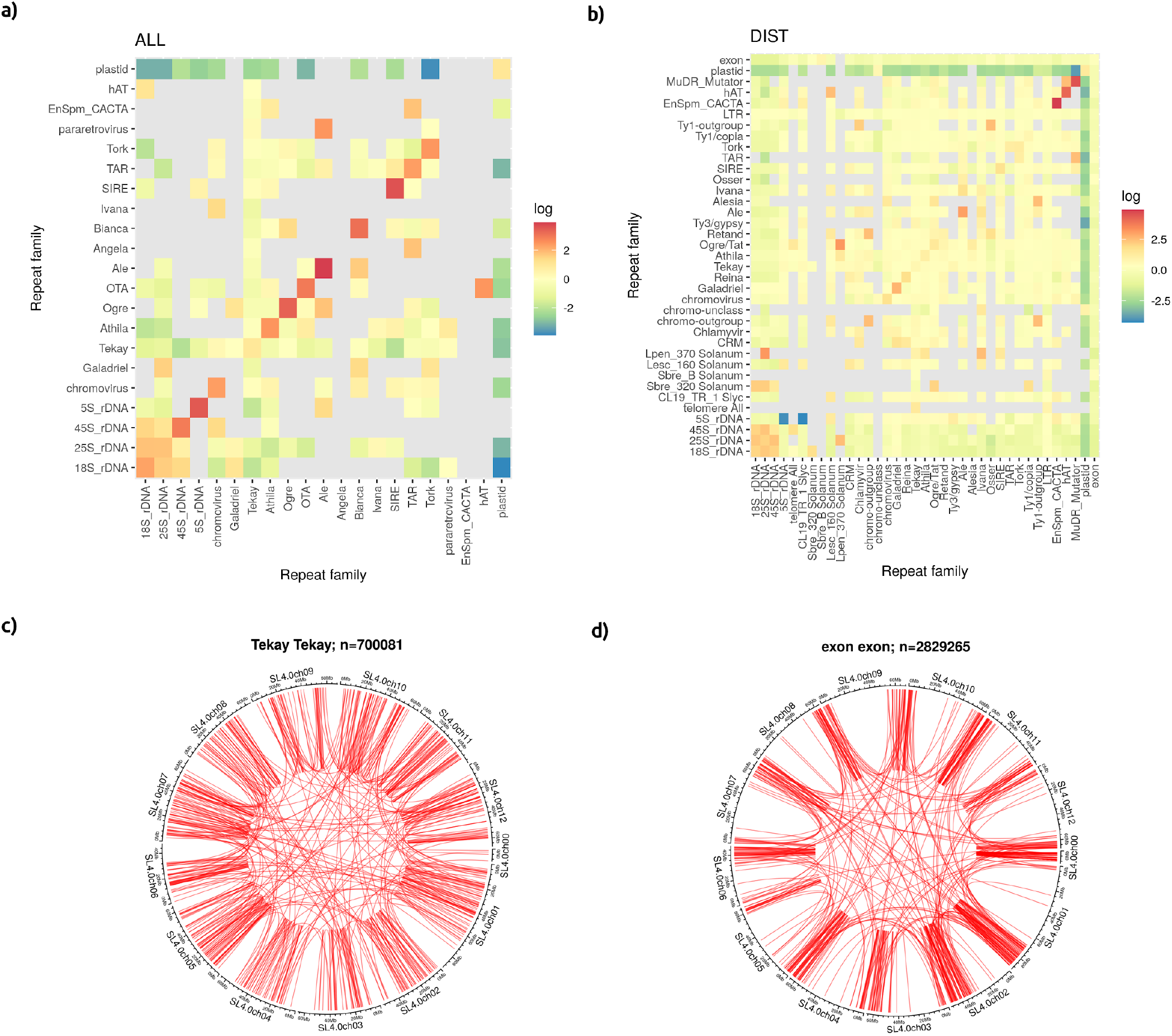
Results of testing the HiC-TE pipeline against 6 sequencing datasets from Dong et al. (2017). Examples from run SRR5748729. a) heatmap of all repeat family contacts based on HiC reads clustered with Repeat Explorer; grey fields are shown for pairs where missing values prevented normalization b) heatmap of long-distance repeat family contacts based on mapping HiC reads to annotated reference genome; grey fields are shown for pairs where missing values prevented normalization c) circular plot of Hi-C-supported interactions between regions annotated as TE family “Tekay”; d) circular plot of Hi-C-supported interactions between regions of the genome annotated as “exon”.

We found that in tomato, interactions among genes are limited to distal arm regions (euchromatin) while repetitive DNA interactions show higher presence in closer proximity to the centromere (heterochromatin) (Fig.2c,d). When two interacting regions belong to the same repeat family we talk about homotypic interactions (diagonals in Fig.2a,b). More than one third of repeat families (rDNA, DNA TEs, Ale, Reina, Galadriel) displayed such a pattern. The homotypic interactions were even more common when short-distance interactions were included, here they formed about 80% of all interactions (Fig.2a, Additional File 5). We showed that a number of individual interacting repeat families displayed a preference for another repeat family when analyzing pairwise long-distance interactions. This was most pronounced in ribosomal DNA, LTR TEs (e.g chromoviruses, Retand, Ogre/Tat, Alesia, Ivana, Chlamyvir, Ty1-outgroup), some tandem repeats (Lpen 370, Lesc 160) and DNA transposons (MuDR Mutator)(Fig.2b).

## Discussion

Here we presented a novel pipeline combining Hi-C data, plant reference genome data and tools for the characterization of genomic repeats. This pipeline can quickly identify and quantify the contacts of specific repeats in the 3D nucleus. Nextflow allowed us to formulate the pipeline in a modular manner and conveniently publish ready-to-run code, be it on individual computers or HPC environments, with docker and singularity containers to support all necessary dependencies and their required configuration. The modules (separate Nextflow processes) communicate via standard file formats.

Therefore it should be straightforward to modify the computation to use a different piece of software or different input data. Compared to the non-repetitive fraction of the genome, for which a plethora of tools and pipelines exist, the repetitive sequences, such as those used in this pipeline, are challenging in terms of reliable mapping and the number that can be successfully clustered into families.

The pipeline contains two modes of repeat annotation, reference-free and reference-based. Being able to compare the results from both increases the robustness of the results. While reference-based data contain chromosomal positions and allow the calculation of distances, the reference-free mode avoids the necessity to discern real and apparent read mapping, which is especially problematic when dealing with repeats and short reads. In the case of tomato analyzed here, we included the unassembled “chromosome 00” in the reference-based mode which should have resulted in a more precise mapping and slightly distorted distance-based calculations, such as binning contacts into TAD and DIST categories.

While we took extra care to provide several modes of normalization to be able to pinpoint statistically and biologically significant contacts among the transposable elements or satellite DNA families, individual evaluation may still be needed in some cases (for more details on normalisation see Methods and Additional File 2). For example, in Fig.2a which shows all contacts that can be attributed to a repeat, results may partly reflect the length of the elements and the ability of Hi-C reads to bridge genomic regions a few hundreds of bases apart. However, the presence of the same result in long-distance contacts (Fig.2b) would suggest that individuals of the respective families tend to cluster and perhaps form exclusive domains in the nucleus. The primary DNA sequences, namely abundant repetitive elements embedded in the genome, may in this way instruct genome folding and aid genome compartmentalization. It is known that different types of chromatin regions tend to fold in different ways, with heterochromatic chromatin displaying a different average Hi-C interaction frequencies compared to euchromatin regions (Homer) [9, 10]. Individual repeats or entire repeat families can play a role in 3D organization by e.g. demarcating TAD boundaries [11] or harboring binding sites for architectural proteins [12]. Our pipeline has a potential, based on frequency of interactions of specific centromeric or telomeric repeats, to reveal these distinct local organizations of chromosomes in interphase nuclei, or even more global ones, such as Rabl, Rosette or Bouquet arrangement [13].

3-D contacts between repeats can possibly participate in processes of gene conversion or ectopic recombination [14]. Gene conversion contributes to LTR retrotransposon homogenization, while ectopic recombination helps to delete genomic regions. Since gene conversion is strongest in ribosomal genes [15], rDNA loci served us as an internal positive control. Indeed, rDNA clusters showed strong interactions with each other (Fig.2a,b). While the high homogeneity of rDNA has a functional consequence (the need for a large amount of the same functional rDNA molecules), the homogenization of TEs by gene conversion could be beneficial in ectopic recombination (and subsequent genome downsizing) and thus represents a tool for the regulation of genome size.

## Methods

Data from six sequencing runs from a Hi-C experiment on the tomato [9], Additional File 5) were fed into our Nextflow [16] computational pipeline “HiC-TE” (Fig.1, Additional File 1). The pipeline combines read trimming with Diachromatic [17], TE annotation with TE-greedy-nester [18] and Repeat Explorer [19], satellite/tandem-repeat annotation with TAREAN [20], TRF [21] and PlantSat [22]. Read mapping is done via Bowtie2 [23] and BBmap with a subsequent overlap/intersection analysis with bedtools [24] and visualization in R/Bioconductor [25, 26] using the following packages: circlize [27], dplyr [28], GenomicRanges [29], ggplot2 [30], gplots [31], karyoplotteR [32], MatrixCorrelation [33], ragg [34], reshape2 [35], Rsamtools [36], rtracklayer [37], stringr [38]. This pipeline generated tables, heatmaps and circular plots showing frequency of Hi-C interaction between repeats and other annotated features in the genome (for code see GitLab repository). Before visualization in heatmaps, the data is normalized to account for the fact that repeat families have varying frequencies. As there are several ways to carry out such normalizations, each with its own biases, we therefore generated heatmaps using a range of normalization techniques (details in Additional File 2 and 5). Joint probability normalization assumes Hi-C contacts occur between independent positions and normalizes contact counts against a product of frequencies of the interacting families. Label permutation uses a sample set with family labels subjected to permutation. Annotation interval reshuffling uses a shuffled version of annotation files to normalize contacts.

## Availability of data and materials

The source code and documentation for the Nextflow pipeline is available at http://gitlab.fi.muni.cz/hic-te/.

## Supporting Information

### Additional Files

**Additional file 1**

Suplementary Figure — Flow chart of the Nextflow HiC-TE pipeline (output from running Nextflow with the -graph switch)

**Additional file 2**

HiC-TE manual

**Additional file 3**

Suplementary Tables — HiC-TE Nextflow pipeline performance on a 4-core 3.0GHz Intel Ubuntu box and in the cloud (MetaCentrum metacentrum.cz). Numerical values are averages of 12 runs excluding TE-greedy-nester reference annotation (is needed only once)

**Additional file 4**

Suplementary Tables — Hi-C tomato (Solanum lycopersicum) leaf mesophyll sequencing runs from project SRP110225 (Dong et al., 2017) used to test the Nextflow pipeline. The individual runs represent different biological and technical replicates (see batch and plant numbers)

**Additional file 5**

PDF files with 6 complete sets of outputs, numbered by SRR ID and the name of output

## Acknowledgements

We thank Christopher Johnson for critical reading of this manuscript. We thank Jan Hoidekr for the help with the deployment of the pipeline in the metacentrum environment. Computational resources were supplied by the project “e-Infrastruktura CZ” (e-INFRA CZ LM2018140) supported by the Ministry of Education, Youth and Sports of the Czech Republic. Monika Cechova is the holder of Martina Roeselova Memorial Fellowships 2020.

## Authors’ contributions

Conceptualization ML, EK, PJ; workflow implementation and data analysis ML, MC, SHN; methodology ML, MC, PJ and VT; docker, singularity container preparation SHN; validation PJ; writing - original draft preparation EK, ML, MC, PJ; writing - review and editing ZK and RH; supervision EK; funding acquisition EK and ML. All authors have read and agreed to the published version of the manuscript.

